# Visually Induced Involuntary Movements

**DOI:** 10.1101/2024.02.06.578999

**Authors:** Alexandra Martin, Joel Ventura, Avijit Bakshi, Sacha Panic, James R. Lackner

**Affiliations:** Ashton Graybiel Spatial Orientation Laboratory, Brandeis University, Waltham, MA 02454

**Author notes:** Correspondence: James R. Lackner, Address: MS 033, Brandeis University, 415 South Street, Waltham, MA 02454, USA.

**Keywords:** Illusory body motion, Involuntary arm, head and torso displacements, Tool use in virtual reality, Joystick control, Spatial disorientation

## Abstract

Looking at a virtual 3D environment with structural features rotating at 60°/s in a head-mounted display soon elicits an illusion of self-rotation and displacement in the opposite direction. We explored in 75 s long trials the effects of visually induced self-rotation on the head, torso, and horizontally extended right arm of standing subjects. The degree of body and limb movement was contingent on whether the arm was extended out freely or pointing at a briefly proprioceptively specified target position, but did not depend on whether the hand held a rod or not. Most subjects in the Free condition showed significant unintentional arm deviations, which averaged approximately 55° in the direction opposite the induced illusory self-motion, and were more than 150° in some cases. In contrast, on average, the deviations in the Pointing condition were a quarter of those in the Free condition. Deviations of head and torso positions also occurred in all conditions. Total arm and head deviations were the sum of deviations of the arm and head with respect to the torso plus deviations of the torso with respect to space. When given a pointing target, subjects were largely able to detect and correct for arm and head deviations with respect to the torso but not for the parts of arm and head deviation that were due to deviations of the torso with respect to space. In all conditions, the arm, head, and torso deviations occurred before subjects began to experience compelling self-rotation and displacement. This is contrasted with the compensations for expected but absent Coriolis forces that are made when stationary subjects make reaching movements to targets during exposure to structured moving visual scenes. These compensations do not occur until subjects experience self-rotation and spatial displacement. These results have implications for vehicle control and maneuvering in environments that induce illusory motion and displacement, and in situations where there is motion in a large area of the visual field. The impact of these effects on joystick control is described and discussed. We also describe the subjective sense of ownership attributed to hand-held objects when experiencing illusory self-motion and displacement.

## INTRODUCTION

On July 20, 1969, during the first manned landing on the moon, Armstrong and Aldrin, while standing, manually controlled the final minutes of the lunar module’s descent to the moon’s surface. The exhaust from the rockets controlling the descent created a large leftward stream of dust that interfered with Armstrong’s ability to see landmarks on the surface of the moon. Aldrin would call out control commands that Armstrong tried to execute. The final landing location was deviated leftward in yaw from that intended. Armstrong and Aldrin in later debriefings reported that the leftward stream of dust and particles likely caused an illusion of rightward displacement of the module that Armstrong tried to correct and that as a consequence his command corrections overcompensated to the left.

This event may represent the most potentially significant instance of spatial disorientation in the history of human flight. In recent years, the use of virtual environment technology in training and in cockpits has greatly increased. Some virtual environments involve visual motion that has the capability of inducing illusory self-motion akin to that reported by Neil Armstrong. We wondered whether such moving visual scenes influence not just apparent self-motion and displacement, but also static and dynamic arm movement control. The Giant Hand Illusion described by Gillingham is one of the most compelling illusions experienced by pilots (Gillingham, 1992). In this Illusion, the pilot experiences a tilt of the aircraft that does not respond to intended joystick commands. It seems as if the joystick commands are opposed by an external force – a Giant Hand – maintaining the unwanted tilt, which if not recovered from could cause a fatal crash.

In earlier work, we found that seated subjects experiencing illusory self-rotation and displacement made reaching errors when aiming at a target, as if they were automatically compensating for the Coriolis forces on the arm that would be generated if they were actually rotating (Cohn et al., 2000). This kind of reaching error is also known as the pseudo-Coriolis effect. We are not consciously aware of this, but during our everyday activities when we are standing and simultaneously turn and reach for objects, substantial Coriolis forces are generated on the reaching arm. This occurs because high peak torso velocities are generated and the peak velocity profiles of the torso and arm overlap nearly perfectly to create a Coriolis force on the arm (Bortolami et al., 2008, Pigeon et al., 2003b, Pigeon et al., 2003a). Such Coriolis forces (CF) are generated when an object moves in relation to a rotating reference frame, CF = −2 m (ω x υ), where m is the mass of the moving object, ω is the angular velocity of the rotating reference frame with respect to an inertial frame, and υ is the velocity of the moving object in relation to the reference frame. During initial exposures to being in a rotating environment, such as a fully-enclosed rotating room turning at constant velocity, an individual’s reaching movements to a target will be deviated by the CF generated, resulting in trajectory curvature and endpoint errors. The individual making a reaching movement will feel him or herself to be physically stationary and it will feel as if an intangible force has deviated the path and endpoint of their reach, and that their arm had not done what was intended. With additional reaches, accuracy will soon be regained and it will then feel as if the CF has gone away; reaches again feel fully normal. When room rotation is stopped and a reaching movement is made, there is a negative aftereffect. It feels as if a CF is again present during the reach, but now that force deviates the arm’s path and endpoint in the direction opposite to the initial errors. This aftereffect is the CNS’s compensation for a now expected but absent CF. CFs are created in everyday life during turn and reach movements (and many other movement patterns as well) and our CNS compensates for them so that we are not even aware of their presence.

We had also found in this earlier work that when seated stationary individuals reach while experiencing illusory self-rotation, they do not make reaching errors representing compensations for CF unless they are concomitantly experiencing self-displacement (Cohn et al., 2000). The sense of spatial self-rotation can be readily elicited by moving patterns of stripes, but an accompanying sense of self-displacement in accord with the velocity of visual movement is more easily evoked by virtual environments with complex structural scenes, e.g., people, buildings. The significance of these results for arm movements in virtual environments is amplified in the Discussion section.

In the experiment to be described, we addressed whether arm control would also be affected by combined illusory self-motion and displacement induced by an HMD showing a rotating 3D rendition of a room with windows and objects and internal structure. We wanted to see whether induced apparent self-motion and displacement would influence the arm when held passively outstretched and horizontal, and when held horizontal attempting to maintain a spatial position that had first been haptically indicated. Conditions with the hand holding a tool, an aluminum rod, were included. The potential influence of eye deviation was also evaluated. In all conditions, the subjects were standing, as Armstrong and Aldrin had been during their lunar landing.

On the basis of pilot observations, we thought that in the tool conditions, the rod would be perceived as an extension of the subject’s arm. We also thought that giving the subject a brief haptically specified target to point to would attenuate arm displacement relative to that associated with the free arm condition. Thus, our guiding hypotheses concerning visually induced involuntary arm displacements were as follows: i) free arm, no tool condition = free arm with tool, ii) pointing arm, no tool condition = pointing arm with tool, and iii) free arm > pointing arm, regardless of tool.

The experimental results below show that exposure to a large area visual flow field that includes self-rotation cues results in major unintentional deviations of the freely voluntarily outstretched arm, with or without the rod, which can reach more than 150°. Intentionally pointing the arm, with or without the rod was also affected, but less so, with the arm exhibiting a nystagmoid pattern with a slow drift of 5-10°, alternating with saccade-like corrections as subjects detected their involuntary arm deviation. Although we were initially interested in visual motion induced deviations or displacements on the arm, we soon found that there were similar effects on the head and torso. We will collectively refer to these visually induced movements as Deviation Effects. We will also compare these Deviation Effects in the period of visual stimulation before versus after the onset of illusory self-rotation and displacement. We will explore whether the perception of self-rotation and displacement is necessary for Deviation Effects to occur, as was the case for the pseudo-Coriolis effect explained above.

## METHODS

Subjects: Twelve members of Brandeis University, both students and staff members, participated after signing an informed consent protocol approved by the Brandeis IRB. They included four females and eight males, ranging in age from 18-65 years. All were in good health and without any physical impairments that could affect their balance control or reaching abilities.

### Apparatus

#### Head Mounted Display (HMD)

Subjects wore an HTC VIVE™ HMD with two OLED eye display units with a resolution of 1080 x 1200 pixels per eye, and a refresh rate of 90 Hz. The binocular field of view was ≍ 110 degrees horizontally and vertically. The HMD presented a 3D visual scene depicting a rectangular room. The room’s inner dimensions were: width 4.7 m, depth 4.7 m, height 3 m. The room had two doors and two windows, and interior decorations including furniture and plants. It was designed to provide a complex, differentiated spatial environment.

When set in rotation and viewed by a standing subject, after a lag, the visual scene invariably would be seen as stationary and the subject would experience compelling self-rotation and self-displacement about their vertical z-axis, which was aligned with the center of the rotating virtual room. The HMD scene rotation rate was set at 60°/sec CW during trials. Pilot tests had shown that CW and CCW rotation induced results that were indistinguishable except for sign.

#### Dual Force Plate

During experimental trials, subjects stood on a dual force plate (AMTI Accusway^TM^) that allowed separate monitoring of the forces and torques exerted by each foot. The force plate was surrounded on three sides by waist-height safety railings that the subject could grasp were it necessary. In addition, one experimenter was always near the subject to provide balance assistance if necessary. The force plate results are described in the online supplement to this paper as they are not directly relevant to the Deviation Effect results, which are the main interest in this paper.

#### VICON Motion Capture System

Eight VICON infrared cameras monitored 14 passive markers that were attached to the subject’s head (HMD), torso, and right arm and hand. The sampling rate was 100 Hz.

#### Tactile target

For trials involving pointing to a haptic target, a tripod was used, with its height set to the height of the subject’s horizontally outstretched arm to give a momentary tactile cue to the subject’s index finger or to the distal end of a hand-held rod. This tactile cue was set to the position where the subject, with their head straight ahead, felt was directly ahead of their right shoulder. It was briefly given while the subject viewed a stationary view of the virtual room in the HMD just before a trial began.

### Experimental Design

The experimental format followed a 3×2×2 factorial design. The trial duration was always 75 s.

#### Independent Variables

The three independent variables were Epoch, Task, and Tool. Epoch had three periods. The first (“Pre”) was the period after trial onset before the subjects’ report of self-motion and displacement. It was calculated as the time from the subjects saying “ready” (to trigger trial onset) to the subjects reporting “moving”. The second period (“Initial”) was set to the length of the Pre period and occurred immediately after it, and represented the initial period of self-rotation and displacement. The third period (“Remainder”) was the time from the end of the Initial period to the end of the trial. Having the Pre period and Initial period of the same duration allowed us to compare them to determine whether visual stimulation prior to the induction of apparent self-motion and displacement could affect arm, head, and torso control.

The second independent variable was Task, with two levels. In the Free condition the subjects only had to hold their extended arm and hand horizontal, and to not correct any arm movements they might sense. In the Pointing condition, the subjects had to hold their arm horizontal, while continuously pointing to a haptically pre-cued location in space for the 75 s duration of each trial. They were instructed to null any arm movements. The two conditions of this variable were counterbalanced for presentation order both within and between subjects.

The third independent variable, Tool, also had two conditions. In the no-rod condition the subjects held nothing in their hand. In the rod condition, the subjects held an aluminum rod (tool) horizontally in alignment with their outstretched arm. The rod was 1.27 cm in diameter, 75 cm long and weighed 270 g. The two conditions for this variable were also counterbalanced for presentation order within and between subjects.

Figure 1A depicts the beginning of a trial in a Free task condition with the rod. Figure 1B shows how the arm and rod were deviated at the end of the Free condition trial. (Figure 1 about here)

**Figure 1.**
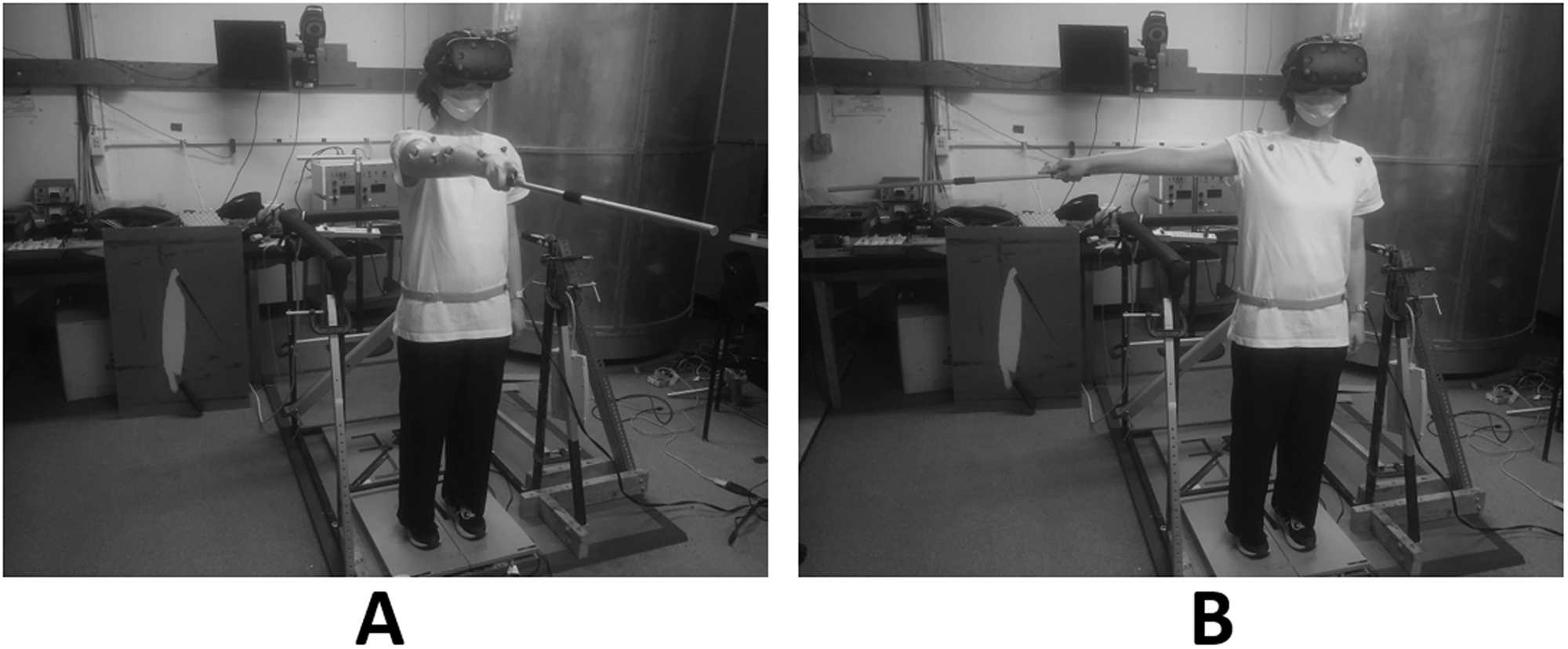
A subject is shown standing on a force plate, wearing an HMD, and pointing forward at the start of a rod condition (**A**), and the azimuthal arm and rod deviation visible near the end of a trial (**B**).

#### Dependent Variables

We collected data for a total of 11 dependent variables related to arm, head, and torso motion. These included the length of the self-motion induction period, six variables to describe the arm, head, and torso positions and average velocities in space, and four variables to describe the positions and average velocities of the arm and head in relation to the torso. The average velocity was computed between the initial forward angular direction and 80% of the maximum arm deviation angle observed in the epoch. We used 80% of the maximum deviation to compute the average velocity because the head, torso, and arm displacements could reach physical limits before a trial was over, and after reaching the maximum deviation (in the Remainder Epoch) there could be zero velocity corruption of the computation of the average velocity of angular movement.

Each subject completed 16 experimental trials that were divided into four blocks, following a 2×2 design for free vs. pointing and no-rod vs. rod conditions. The four possible combinations of Task and Tool were presented four times for each subject, and the order of presentation was counterbalanced across subjects. There were four counterbalanced experimental orders and each subject was assigned randomly to one. Three eye-deviation control trials described below were also included.

In the No-Rod Pointing Condition, the position of the pointing target was established at the start of the trial by having the subject place their index finger on the top of a tripod at shoulder height straight ahead of their right shoulder. In the Rod Pointing condition, the distal tip of the rod was placed in contact with the top of the tripod before the trial started. The tripod was removed at the start of the trial. In the Free Arm Condition, the subject had to raise their right hand and arm to the straight-ahead horizontal position, but there was no tactile target or requirement to point to or maintain their arm in the straight-ahead position, just to hold it horizontal. In the Free-Rod Condition, they held the Rod horizontal with their outstretched arm, but were instructed not to correct any deviations during the trial.

#### Eye Deviation Control

Viewing a rotating visual scene will elicit an optokinetic nystagmus with slow phase compensatory for the direction of the scene motion and, as a consequence, the average overall position of the eyes will be deviated relative to the resting position. A free no-rod control condition was added to identify any effect of eye deviation per se. The subject held their hand in a horizontal straight-ahead position, closed their eyes and then voluntarily deviated them fully to the right for 75 sec. Three eye-deviation trials were included with one before the first experimental trial, one midway (after completion of two four-trial blocks), and one after the last experimental block.

#### Preliminary instruction/demonstration/test

Before the start of the experiment, subjects underwent pointing versus free arm demonstrations to ensure that they understood the different tasks. They first were asked to assume they were in the pointing condition, and to hold their arm horizontally as if pointing to the remembered tactile target. The experimenter then physically pushed the subject’s arm left and right 15 to 20 degrees in the horizontal plane. If the subject did not immediately correct their arm deviation, the experimenter repeated the demonstration test until the person understood that they needed to keep pointing at a specific location in space during the whole duration of a trial. Next, the subjects were told to assume they were in the free-arm condition of neither pointing nor correcting. The experimenter then again moved the subject’s arm in the horizontal plane and if a subject resisted their arm being pushed or immediately returned it to the straight ahead (“correcting”), the experimenter repeated the instructions and the test until the subject understood not to correct arm deviations in this condition. These simple demonstration tests were important, because we had found in pilot testing that the subjects had no problems with the pointing task, but even though we had explained in words how the free arm task was different, some subjects still returned their hand to the felt straight ahead position when they detected an arm deviation. These simple demonstrations made the “do not correct” distinction clear to all subjects.

#### Experimental Procedure

After the VICON markers were attached to the subjects, instructions were given followed by the Pointing and Free Arm Condition demonstrations described above. The subjects then took their position on the force plate and put on the HMD, with its visual scene stationary. Before the beginning of each trial, subjects were asked to close their eyes, hold their right arm horizontally (with or without a rod), and were told trial type – free arm or pointing – and to report out loud “moving” when they felt compelling self-motion and displacement. The experimenter started the virtual room rotation and signaled verbally to the subject that the experiment was ready to start. The subject then opened their eyes while simultaneously saying “ready” to the experimenter who started a stopwatch. The experimenter stopped the stopwatch when the subject reported “moving” and recorded the onset time of experienced self-motion.

For the pointing trials, the subject was given a tactile cue by placing their index finger or the distal end of the rod on the top of the tripod. As soon as the subject said “ready” while opening their eyes, a second experimenter removed the tripod. For the free blocks, subjects were instructed to hold their arm horizontally without trying to maintain it in a particular position.

After each trial, the second experimenter checked that the subject’s feet were still in the proper position and corrected head and torso positions if they had changed during the trial. Subjects were given optional breaks after completion of a four-trial block (rarely chosen) and a mandatory five-minute break halfway (two blocks) through the experiment, during which they sat down.

#### Statistics

To keep the total number of variables equal to or less than 10 in a given MANOVA, as recommended by Stevens (1980) and Field (2009), we divided the dependent performance variables into two areas of interest:

1. AHT re Space-- Movements of the arm, head, and torso with respect to space (six variables).
2. AH re Torso-- Movements of the arm and head with respect to the torso (four variables).

We performed two 3-way MANOVAs, one for each group of variables in the two areas of interest. The MANOVAs tested for main effects of the independent variables 1. Epoch with three levels (Pre, Initial, Remainder), 2. Task with two levels (Free, Pointing), and 3. Tool with two levels (rod, no-rod). We also tested for significance of all possible interactions among the three independent variables. Table 1 shows the full testing structure in detail.

**Table 1.**
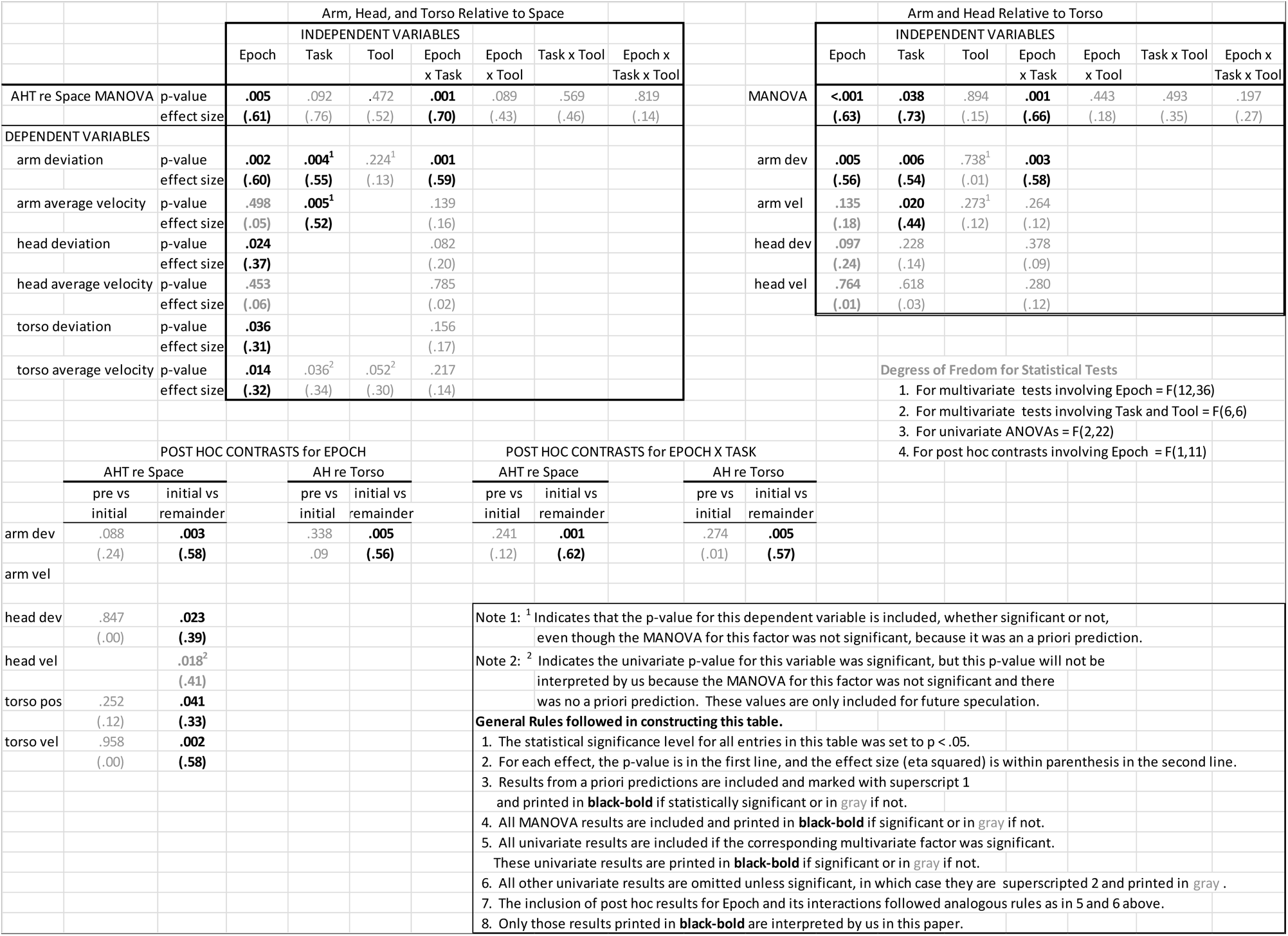
Summary of Study Main Statistical Results.

Roy’s Largest Root (Θ) statistic was used to determine the significance of each MANOVA. It is most powerful when differences are due to the first variant (Field, 2009, Stevens, 1980). We tested the assumption of sphericity for each MANOVA. If it was violated, we adjusted the significance level of the follow-up ANOVAs using the Greenhouse-Geisser correction. This correction is only necessary for tests involving the main effects of Epoch or its interactions, because this was the only independent variable with more than two levels. In addition, if there were significant main effects or interactions in the Epoch ANOVA, we performed post hoc tests to determine which differences among the three levels of Epoch (Pre, Initial, Remainder) were significant. Finally, we did a separate ANOVA on the induction period (Pre) as a function of Task and Tool.

#### Summary Statistical Comparisons

The statistical results are summarized in Table 1 for the reader’s guidance through the complex pattern of findings. Table 1 includes a detailed explanation of the coding used and rules followed in its construction. The significance levels are shown as well as the Partial Eta Squared (η^2^) effect sizes.

To control the overall Type I error rate in interpreting the results we first checked the multivariate significance of each factor, and when significant, checked the ANOVA for each dependent variable to determine which ones contributed to the multivariate significance. The Epoch factor has three levels and we further checked the post hoc contrasts to see which differences between levels caused the significant ANOVA. Only when we had a priori predictions (superscripted 1 in Table 1) did we test for an effect whether it was significant at the previous level or not.

## RESULTS

The mean onset time for illusory self-motion and displacement (induction period) across all subjects and conditions was 10 ± 7.7 s (mean ± std dev). The times for onset were 10.2 ± 7.6 s for the *Free – no rod* condition; 9.3 ± 6.5 s for the *Free - rod condition*; 10.3 ± 7.4 for *pointing with no rod*; and 10.2 ± 9.3 s for *pointing with the rod*. An ANOVA showed that neither Task nor Tool had a significant effect on the length of the induction period. (F(1,11) = 1.4, p >.25 for Task type; F(1,11)=.1, p>.75 for Tool factor).

The main analyses of angular deviation magnitude and angular deviation velocity were done in two different coordinate systems. The first was a system fixed in inertial space, and considering that the head and arm are attached to the torso so that when the torso rotates it carries the arm and head along as well, the second coordinate system was fixed to the torso so that we could measure arm and head movements with respect to the torso.

### Angular Deviations

#### Tool Effects

Table 1 shows that there was no significant effect of Tool in either the multivariate or in any of the univariate dependent measure ANOVAs in this experiment, either re space or torso. Therefore, to simplify Figures 2 and 3 below, the data from the Rod and No-Rod Tool conditions were averaged.

**Figure 2.**
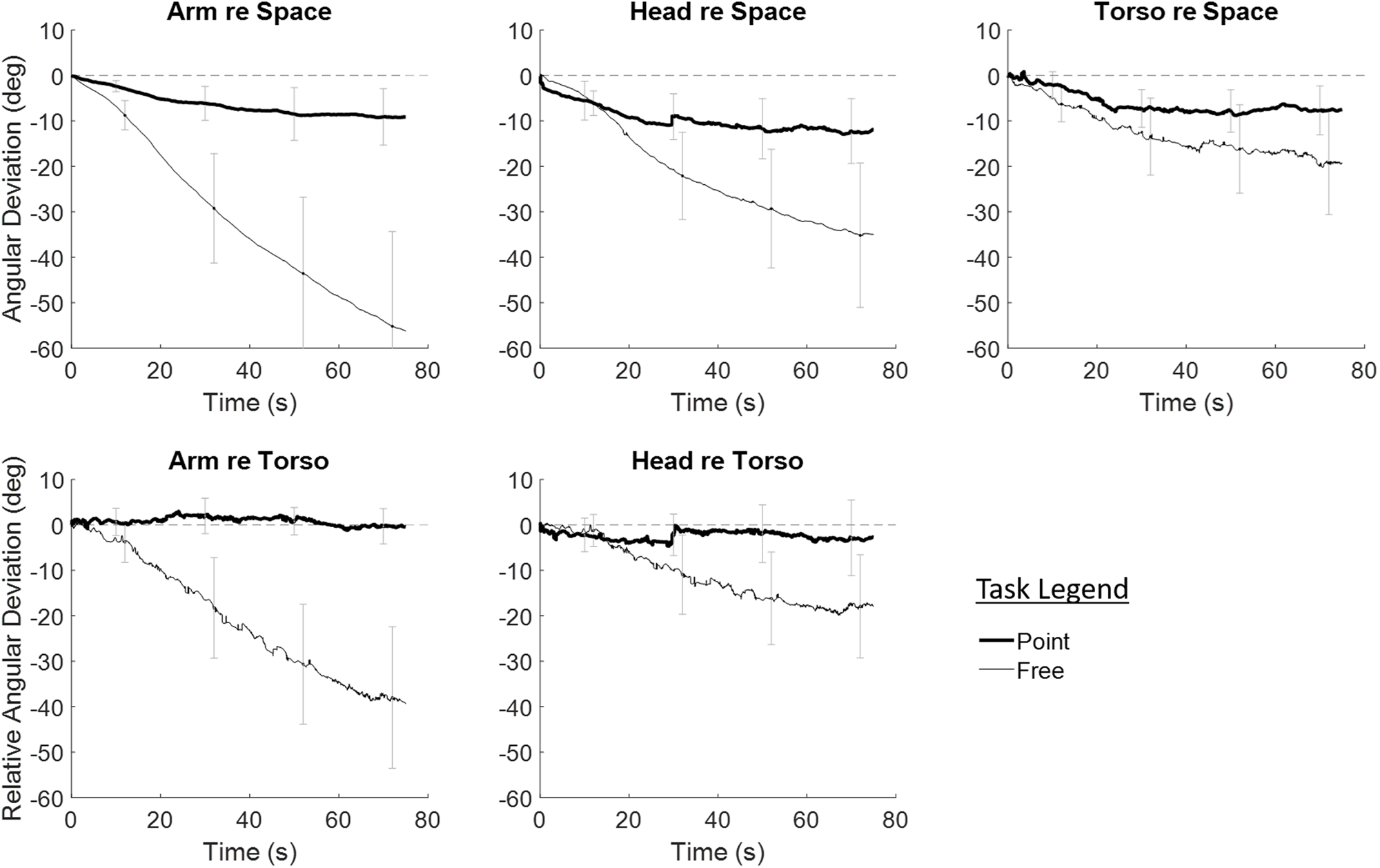
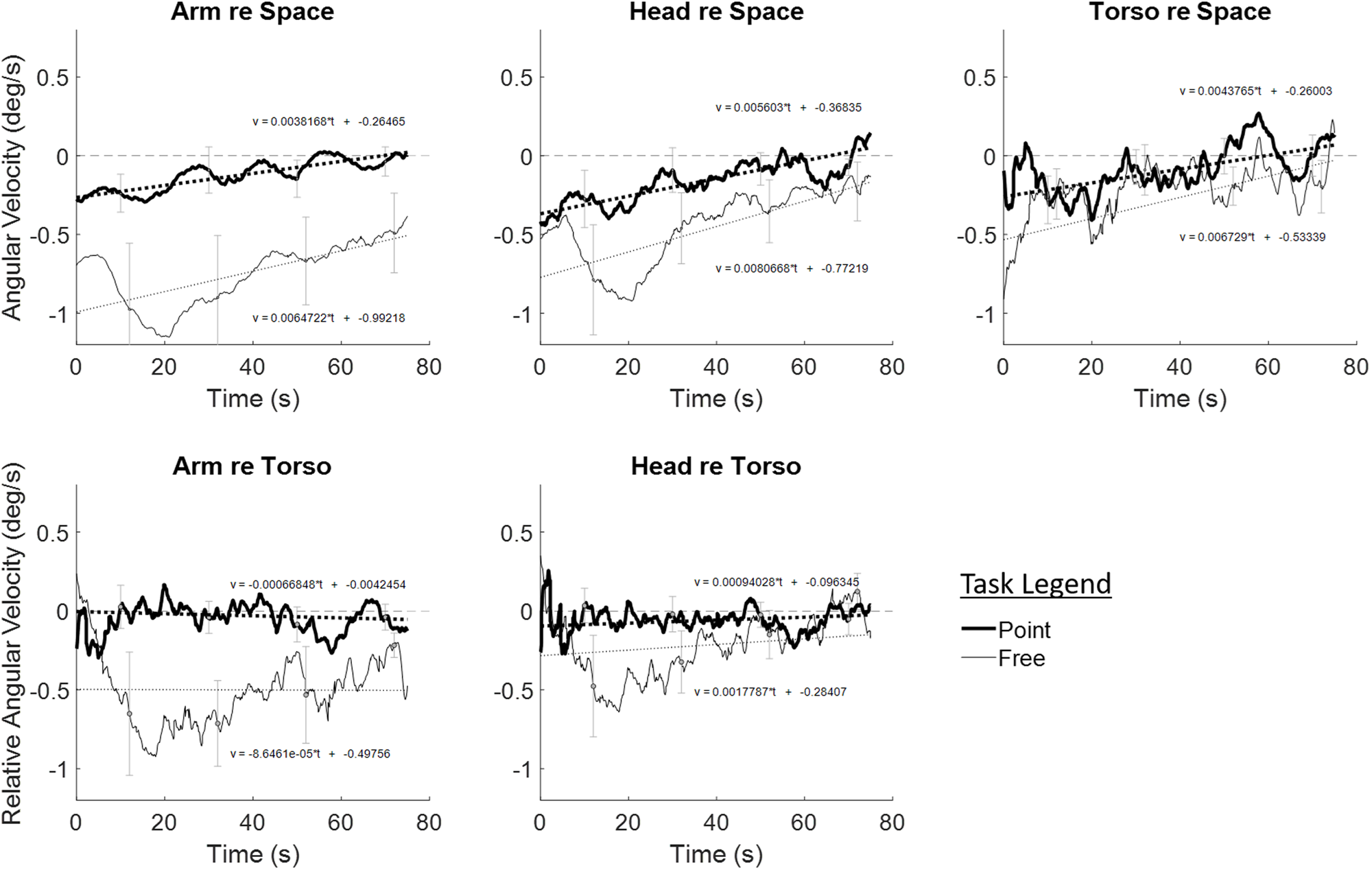
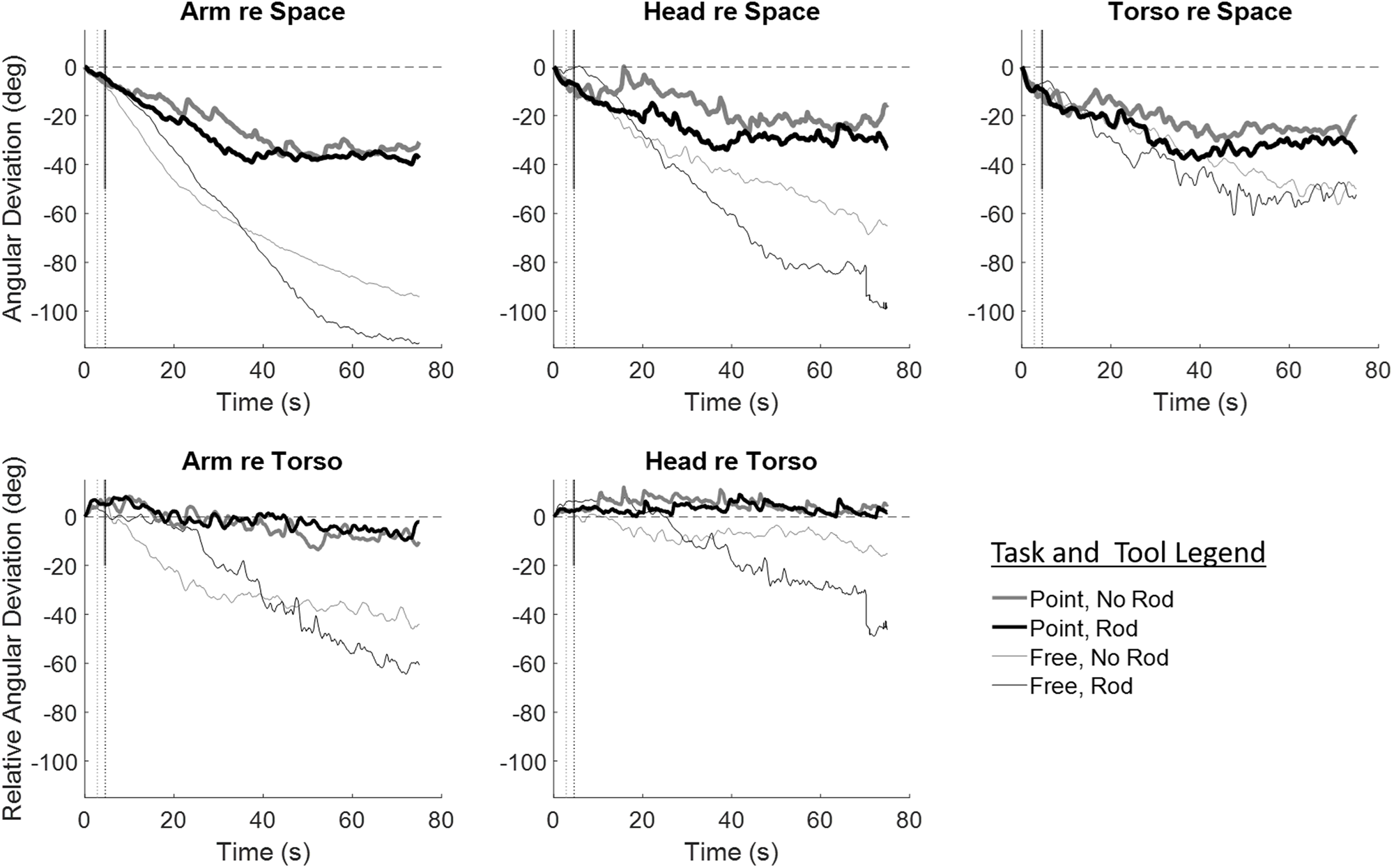
The subject averaged azimuthal angular deviation for arm, head and torso in space (upper row) as a function of time. The mean deviation during the Free task was negative (CW, opposite to the CCW illusion of self-rotation), with the magnitude of deviation in the order A>H>T. For the Pointing task, the deviations were relatively small, nearly equal in magnitude (CW as well). Standard deviation error bars are shown at selected time points. The deviations of the arm and head with respect to the Torso are shown in the bottom row. In the Free task condition the arm moves significantly beyond the torso (more than double relative to Torso and more than three times in space). The presence and absence of tool conditions have similar deviations for the arm. The head moves as much relative to torso as torso moves in space. In the Pointing condition, the arm and head have minimal deviation relative to torso motion, indicating that they are on average primarily moved in space by the torso except in the cases of extreme arm deviations. Standard deviation error bars are shown for selected time points.

**Figure 3.**
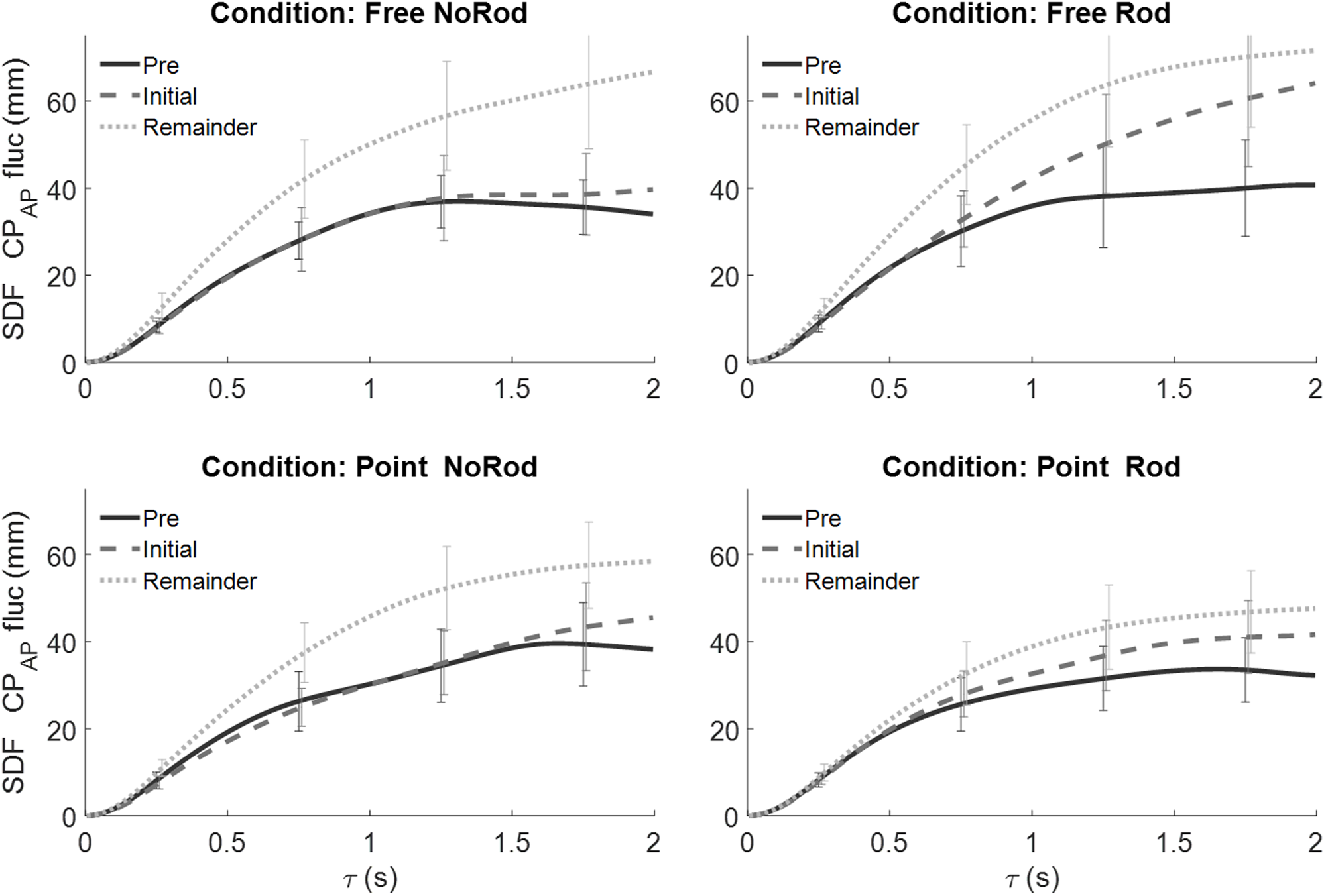
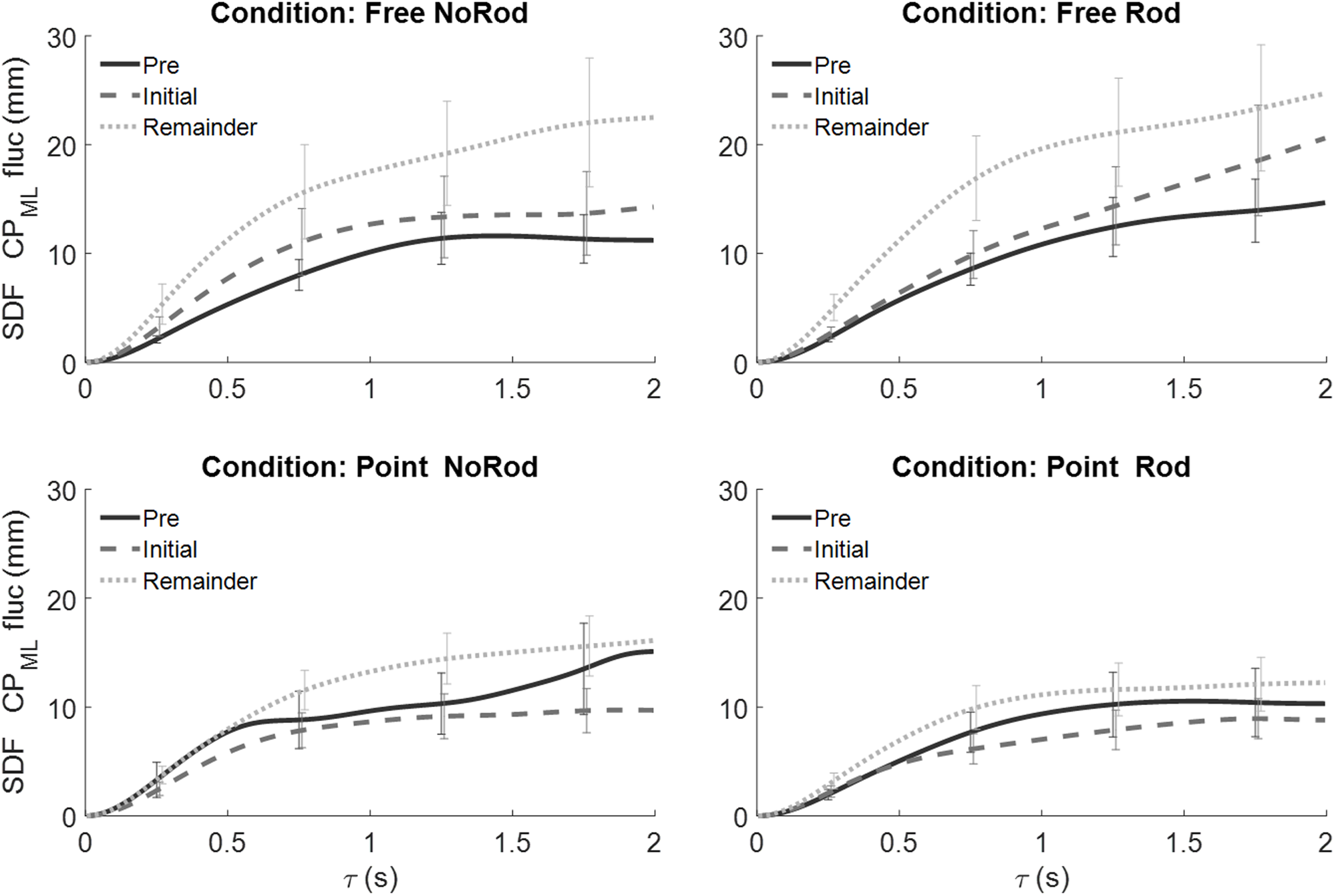
The average angular speeds of arm, head and torso deviations as a function of time for the Free and Pointing conditions. The velocity during Free has a larger magnitude than during Pointing. The linear best fits (dotted lines) suggest that the arm, head and torso velocity re space have a positive slope (shown in the top panel) as a function of time, and the magnitudes reduce to near zero by the end of the trial. The velocities re Torso (bottom panel) show that the arm and head were moving along with the torso (zero relative speed) in the Pointing condition.

We thought a Tool effect was a real possibility, because although the rod only weighed 270 g compared to about 4.6 kg for the average human arm, the center of mass of that rod was approximately 35 cm beyond the end of the outstretched arm during the experiment, and that would increase the moment of inertia of the average male arm by approximately 20%, and by about 30% for the average female arm. These are significant changes, yet they did not significantly affect the deviation motion. This result shows that the motor control system does not apply a constant torque on the arm as a result of the rotating HMD visual stimulus, but instead treats the arm plus rod as a new single unit, maintaining similar angular rates of motion. (Figure 2 about here)

#### Task Effects

The arm, head, and torso angular deviations relative to space and re torso as functions of time are shown in Figure 2. The analogous curves for deviation velocities are shown in Figure 3, and the statistical results are shown in Table 1. The Task main effect was not significant when all arm, head, and torso variables were analyzed in the same MANOVA (p =. 09). However, the effects of Task on the arm variables were still analyzed, because this was a planned comparison based on pilot tests conducted prior to this experiment. The largest effect shown in Figure 2 is the effect of Task on arm deviation. Re space the maximum deviation of the arm in the Free condition is more than four times that in the Pointing condition, −56° vs. −9°, and this difference is similar re torso, −39° vs +3°. Table 1 shows that these differences were statistically significant and that the effect sizes were also large. Angular deviation is also greater for the head (−35°) and torso (−19°) in the Free condition vs. the Pointing condition (−12° and −7.5°), but these differences do not reach statistical significance. The standard deviations are also greater in the Free condition than in the Pointing condition (p < .001 re space, p= .004 re torso; tested by Wilcoxon Signed Rank test at eight pairs of points spaced at equal time intervals in trials). The Task effect occurred because the subjects detected the arm deviations in the Pointing condition and attempted to correct them, which reduced the arm deviations and average velocities compared to the Free conditions. These corrective movements will be described in more detail below. Examination of the subplots in Figure 2 clearly show that the Free arm condition deviations relative to space are due to the sum of the torso deviations re space −19° plus the deviations of the arm with respect to the torso −39°. In the re space analysis, the deviations of the arm and head are both statistically significant, but in the re torso analysis only the arm deviations are significant. This indicates that the head moved almost fixed with the torso, while the arm deviations were not solely attributable to torso deviations. The remainder of the effects in the re torso analysis are similar to those in the re space analysis.

In all conditions, the average torso movements in space were in the opposite direction to apparent self-motion, and the arm and head deviated along with the torso but then the arm continued to deviate even further with respect to the torso. Subjects in the Free and Pointing conditions sometimes reported that their arm seemed to “take on a life of its own”. Several subjects in the Free-rod condition reported that it sometimes felt as if the rod were pulling their arm farther and farther laterally to the side, and that the rod and their arm could continue to feel in motion with respect to their torso but without deviating further. Subjects in both the Free-rod and Point-rod conditions did not report a change in the rod angle with respect to the arm, the rod and arm always moved as a block, and this actually was the case as shown by the video recordings.

Another key result from the Pointing conditions is apparent in Figure 2. The arm and head deviations re torso in the Pointing condition hover around zero, but re space they hover around the torso deviation re space. For example, the average difference between the arm deviation re Torso drops from 21 degrees in the Free condition to less than one degree in the pointing condition. The Arm and Torso deviations re space both remain at approximately −6.1 degrees in the pointing condition. **This means that when subjects are given the Pointing instruction they were able to detect and correct their arm deviations with respect to their torsos, but they were not able to detect and correct that part of their arm deviations that resulted from the deviation of the torso with respect to space.** This conclusion is supported by the significant task effect in the re Torso MANOVA.

#### Epoch Effects

Table 1 shows that the only significant univariate velocity effect for Epoch was for torso velocity. The post hoc contrasts showed that that difference only occurred between the Initial and Remainder Epochs. There were no significant differences between the Pre and Initial Epochs. This has important implications for the conditions under which these deviation effects occur and will be addressed in the discussion.

Re space for both Free conditions, the arm deviation increased from about 6° in Pre to 51° during the Remainder Epoch, while in the Pointing conditions the arm deviation increased from ≈ 3° in Pre to about 6° during Remainder. The head and torso had smaller deviations. The head deviation, during the Free condition, increased from 6° during Pre to about 27° during Remainder. The torso deviation during the Free condition increased from 3° in Pre to about 18° during Remainder. The head and torso deviation changes during the Pointing task from Pre to Initial were small, ≈ 5° from visual motion onset and ≈ 5° during the Initial period.

#### Factor Interactions

There was a significant interaction between Epoch and Task for arm deviation, and the post hoc tests in Table 1 show that this was due to the difference between the Initial and Remainder Epochs. This occurred because in the Free condition the subjects’ arms continued to drift without correction, while in the Pointing condition the subjects attempted to correct this drift when they detected it. It took some time for that drift to be detected, consequently it was more likely to be detected and corrected in the Remainder Epoch, which lasted much longer. There were no other significant interactions in deviation magnitude or velocity in these MANOVAs.

### Angular Velocity

Taking the derivative of the deviation data to obtain velocities is a noise inducing process. To see the long-term trends in velocity, we filtered the velocity data with a rolling average filter, also called a sliding window average, which is a low-pass filter. We used a window width of 7.5 seconds, which is 10% of the total trial duration. We also plotted the linear fits to the unfiltered data for each curve condition in the graphs and plotted the standard deviations of the unfiltered data as vertical bars approximately every 20 seconds. The result is shown in Figure 3.

#### Tool Effects

As was the case when we looked at angular deviations, Table 1 again shows that there was no significant effect of Tool on deviation velocities in either the re space or re torso MANOVAs.

#### Task Effects

Table 1 shows that the only significant Task deviation velocity result was for arm velocity, and this was true for both the re space and the re torso MANOVAs. The linear fits to the velocity data for the arm and head show large differences from zero for both slope and intercept in the re space analysis but not in the re torso analysis. The average velocities of the arm and head re torso in the pointing condition are both approximately .05 deg/s. **This supports the finding from the angular deviation results that the subjects cannot, or at least do not, correct their deviation errors re space in the Pointing conditions, and only correct errors re torso**.

#### Epoch Effects

As shown in Table 1, the only significant effect of Epoch on the deviation velocity occurs for the torso velocity re space. Figure 3 shows that re space, for the arm and head in the Pointing condition, the maximum deviation velocity occurs in the first few seconds and then gradually decays approximately linearly to zero over the course of the trial. The slopes of those linear fits are significantly different from zero in the re space subplots, but the slopes are not different from zero re torso, again confirming that the subjects correct their re torso deviations, but not their re space deviations. For the analogous Free conditions, it appears to take at least 10 times longer (about 20 seconds) to reach the maximum deviation velocity before the approximate linear decay towards zero begins.

Note that Table 1 shows that Epoch affected the angular deviations of the arm and head, but not their angular deviation velocities. Normally deviation magnitude and velocity results should be highly correlated. However, the duration of the Pre- and Initial Epochs was much shorter than that of the Remainder Epoch, so the same velocities could result in very different angular deviations in the different Epochs, and this can break the correlation.

#### Centripetal force

We wondered whether subjects experiencing Deviation Effects about their own vertical Z-axis while holding the rod (which seemed to move with their arm as a single unit) would experience a “centrifugal force” on their hand. We calculated the centripetal force (F_CP_) that would have to be exerted by the hand to hold the rod in position during actual ω=60°/sec rotation, F_CP_ = mass_rod_ * ω^2^*(Length_arm_ + Length_rod_/2). The value is only about 0.31 N (0.12 times acceleration due to gravity ≈ 32 grams) and is likely near or above the detection threshold, albeit not by much. On questioning, subjects reported that they had not felt any radial centrifugal force on their hand from the rod.

#### Nystagmoidal Arm Deviation Reversals

Although the average values for the pointing movements of the arm are close to the initial start position, the arm and torso actually exhibit nystagmoid-like saw tooth patterns where the arm at first lags the apparent direction of illusory body motion. Then, either it or the torso makes a jerk-like motion to reverse the arm displacement motion. Some subjects reported on questioning post-experiment that it felt as if their arms “drifted” without their awareness and then reached a detectible displacement that they corrected to keep the arm on the remembered target position. The pattern of single trial plots for one subject with large amplitude reversals is shown in Figure 4. The differences in nystagmoid-like responses between the Pointing and Free conditions are easily seen re Space (top row), but are less obvious re Torso (bottom row).

**Figure 4.**
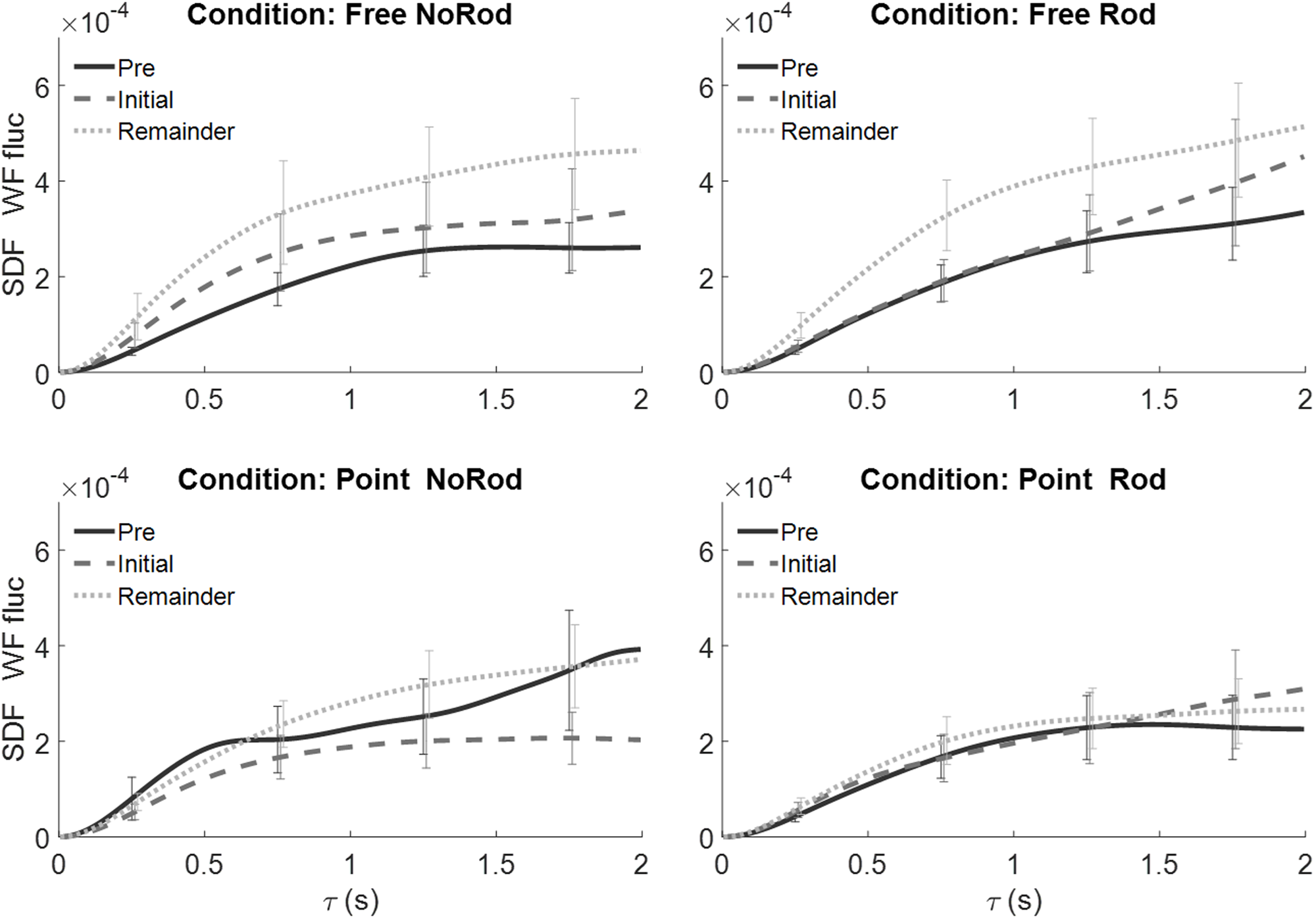
The nystagmus like motion reversals for the Arm, Head, and Torso deviations in space (top row) and re torso (bottom row), of a single subject for two distinct trials - one of Pointing without rod, one Pointing with rod, and two for Free with and without rod. The vertical lines are the motion onset times (there are four lines: two shorter solid lines for the Pointing trials, and two longer dotted lines for the Free trials. Three of the vertical lines are too close to distinguish, as the onset times are nearly identical between 4.3 to 4.6s). Some subjects had prominent body motion nystagmus, while others scarcely showed it. Note that this subject showed much larger than average deviations.

To quantify these visual differences, we created two new measures. When subjects were exposed to the rotating visual environment, their arms drifted predominantly in one direction, but periodically this direction of movement temporarily reversed; how often this reversal occurred in a given trial was called the Number of Reversals (NOR). The percent of trial time the arm or other body part was moving in the reversed direction was the second measure, Duration of Reversals (DOR). We then reanalyzed the arm and head deviation data over entire trials with a MANOVA in which the independent variables were the Coordinate System (re Space or re Torso), Task, and Tool, while the dependent variables were NOR and DOR. There were significant main effects in this MANOVA for Coordinate System (Θ=4.94, F(4,8)=8.0, p<.003, η^2^=.83) and Task (Θ=2.12, F(4,8)=4.39, p=.036, η^2^=.69). Subsequent ANOVAs showed that the significant multivariant Coordinate System effect had contributions from Arm-NOR (p=.008), Arm-DOR (p<.001), Head-NOR (p=.005), and Head-DOR (p=.006). The significant multivariate Task effect had contributions from Arm-NOR (p=.004), Arm-DOR (p<.001), and Head-NOR (p=.029), but not from Head-DOR (p=.111). In summary, these results combined with the marginal means show that the reversal effects are more frequent and of greater duration in the Pointing conditions than in the Free, and higher frequency components are being added to the arm and head re torso by the torso movements.

#### Eye Deviation Control

We analyzed the eye deviation control data with a one sample MANOVA test (a multivariate analog of the univariate one sample t-test) to determine whether eye deviation alone produced effects different from zero on our six measures of arm, head, and torso deviations. The MANOVA result was not significant (Θ=.870, F(5,7)=1.218 p =.391). Since this was a control condition, we wanted to ensure that we could rule out eye deviation as a cause of the Deviation Effects, so we also examined the six univariate ANOVAs. This is tantamount to doing six two-tailed single sample t-tests, testing whether each one of our six dependent variables differed from zero during the eye deviation control condition. For the arm and torso position and velocity, the p-values were all greater than .852. But for head position p = .026, and for head velocity p = .035. These results indicate that while the eye deviation control had no significant effect on the arm or torso, it did induce head deviations in the same direction as the eye deviations, rightward. These head deviations may have been due to an artifact of eye movement discomfort, as described in Appendix 1.

### Are Self-Motion and Displacement Necessary for Deviation Effects to Occur?

One reason we divided the experimental trials into three Epochs (Pre, Initial, and Remainder) was to see whether self-motion and displacement were necessary for the Deviation Effects to occur. This is why the Pre and Initial durations are of the same length. If apparent self-motion and displacement were necessary for Deviation Effects to occur, then no arm, head or torso deviations should occur until self-motion is reported. Therefore, there should be significant deviation differences between the Pre and Initial Epochs, and as the post hoc tests in Table 1 show, that was not found. For most subjects, self-motion occurred around 10 seconds into the trial. Figure 2, shows clear evidence of arm, head, and torso deviations that return almost linearly to the beginning of the trial. To quantify these observations, we did one sample one tailed t-tests to determine whether there were significant deviations in the Pre Epoch, and that was true in every case (p < .05 in all tests). These results again confirm that Deviation Effects begin in the Pre-Epoch before self-motion and displacement are perceived. Although self-motion and displacement are not necessary for Deviation Effects to occur, they may increase the magnitude of deviations. More than half of the individual plots of position versus time show increased deviation velocity when the subject first reports self-motion. However, this effect is small and variable and as shown by the lack of a difference in velocity between the Pre and Initial Epochs in Table 1, it does not approach statistical significance.

#### Analysis of balance

A Stabilogram Diffusion Function (SDF) analysis of the force plate results and their relevance for the present findings are presented in the online supplementary material.

## DISCUSSION

We have found how whole-field visual motion affects the perception and control of arm, head, and torso motion and how it is influenced by Task goals (Pointing vs Free arm), and how it is little influenced by holding a Tool in the hand. Subjects, at the start of a trial, initially sense themselves stationary, immersed in a CW rotating virtual environment. For most subjects, the arm, head, and torso Deviation Effects described above begin in a few seconds and the rate of deviation quickly maximizes and then slowly decreases over the remaining duration of the trial. In less than 10 sec, on average, they experience themselves rotating and displacing CCW in relation to a stationary background. As was described in the Introduction, the perception of self-rotation and displacement is necessary for compensations for expected Coriolis forces to occur. But we have found that perceived self-displacement and rotation are not necessary for the Deviation Effects to occur. This means that Deviation Effects can occur in a much wider range of conditions involving moving visual scenes and therefore can be a more common problem.

Subjects in the Pointing conditions, when given a proprioceptively cued target in space to point to, still exhibited Deviation Effects but to a lesser extent than in the Free condition. Their arms, heads, and torsos slowly drifted without their awareness in the direction opposite that of induced motion, but at a rate that was below a velocity detection threshold. For the arm, its total drift with respect to space is the sum of its deviation with respect to the torso plus the deviation of the torso with respect to space. When the arm exceeds a position deviation threshold, the subjects realize that a deviation has occurred and they make a rapid corrective movement. This process continues over time, and the subjects continue to make a series of nystagmus-like movements. However, these corrective movements only correct the component of arm deviation that is due to the drift with respect to the torso. The component of arm drift due to torso drift with respect to space remains (more details in Appendix 4). The result is that the arm continues to drift with respect to space in the Pointing conditions, but at a lower average rate than in the Free conditions. These results indicate that we could eliminate or at least minimize the arm deviation re space if we stabilized or fixed the torso in space.

There are significant Task effects for arm position and velocity in the re space results in Table 1, and at first that would suggest the motor control system can correct for passive deviations re space for the arm. However, when one compares the arm Task effects re space with those re torso in Table 1, it is clear that these effects are statistically almost identical. That means the arm re space effect only occurs because the motor control system is controlling the arm relative to the torso. If the motor control system were using additional information from the arm relative to space, then that would add more systematic variance, and the significance of the Task effect for arm would be much greater in the re space ANOVAs than in the re torso ANOVAs, and that is not the case.

Subjects could not detect passive deviations re space. At the end of some blocks we had told subjects to freeze in position, and then removed their HMDs. Most subjects had detected some arm deviation, but were surprised at how far their arm had actually moved. This is what one would expect if the subject could detect the re torso deviation but not the total re space component. Furthermore, most were shocked that there was any head or torso deviation at all. This inability to correct for arm and head errors re space can also be seen for one subject in Figure 4. Note that in the re torso coordinate system going from the Free to the Pointing conditions reduces the errors to near zero, but in the re space coordinate system going from the Free to the Pointing conditions only reduces the errors to the level of the torso re space. In other words, the arm and head errors asymptote at the level of the torso, and do not approach zero.

Compared to the Free conditions, the arm, head and torso deviations were much smaller during the pointing conditions. Both total azimuthal angular deviation and the average velocity were significantly smaller for pointing versus free conditions. Most subjects showed small arm pointing errors. Some were unaware of these pointing deviations and reported that their arm did not move at all. Others would become aware of the mis-pointing and make intermittent corrections to the initial pointing direction straight ahead. Subjects who corrected their errors would initially show an arm deviation at a rate below their velocity threshold of awareness, and then, when it reached a positional error threshold, they would correct it back in a “nystagmus like” way. All but one subject made error deviations in the CW direction (opposite to the direction of their self-motion perception). The single subject who made CCW errors was unaware of any arm deviation.

We were curious to ascertain whether our hypothesis that subjects holding a rod would internalize it as an extension of their arm rather than treat it as an external object. The subjects all felt the rod to move with the arm as a spatial unit. We found no differences between holding or not-holding a rod within the Free and the Pointing conditions indicating that subjects internalized the hand-held rod as part of their own body (extended arm) and showed the same effects from induced self-motion as with just their arm. The tool was felt to move with the participant during apparent self-motion and displacement as if it were a part of their body.

In our experiment, we generated an illusory sense of self-rotation and displacement through the visual channel. We found this was sufficient to generate non-volitional limb movements (not just an illusion of limb movement) as well as torso and head movements. In most subjects it overrode proprioceptive cues signaling that the arm, head, and torso were deviating by significant magnitudes. These results show that whole field moving visual stimuli can manipulate our body’s estimations of its dynamic configuration. This has relevance for training paradigms using VR environments. In our experiment, no virtual limb representations were presented in the virtual environment, and thus no artificial manipulation of the sense of limb configuration was used. We found that it is possible to induce body movements using visual stimulation that are not always detectable by the body’s proprioceptive apparatus. Subjects can underestimate the extent of arm, head, and torso displacements, or not even detect them.

An important feature of the experimental findings is that the deviations of the arm, head, and torso in all conditions regardless of task or tool were already present in the Pre-period before the onset of apparent self-rotation and displacement. This surprised us because in our earlier work on pointing movements to targets during exposure to whole field visual motion compensations for “expected” Coriolis forces did not occur unless a sense of both self-motion and self-spatial displacement were present (Cohn et al., 2000). That study did not include reaches in the period before the onset of self-rotation and displacement. Displays that had evoked only a sense of self-rotation but without displacement had been ineffective.

In the original Cohn et al study, subjects experiencing CCW rotation and displacement had made leftward reaching errors. In the present study, subjects experienced CCW self-rotation and displacement and were showing right-ward (clockwise) arm, head, and torso deviations both before and after the onset of self-motion, e.g. during Pre, Initial, and Remainder periods.

The difference in direction for the two paradigms suggests that two separate processes are occurring. To test whether this was possible, we had ten new individuals stand on the dual force plate while wearing the HMD developed for the current study. While standing upright their task was to raise both extended arms up and down in parallel rhythmically through an angle of about 150° from thighs to head level. They began the movements with the display stationary and continued the movements when visual motion was initiated. They reported when they perceived self-motion and deviation of their moving arms. As the arms were raised they deviated leftward and no subjects felt or made the deviations until they experienced self-motion and displacement. The leftward deviation was virtually absent after 20 cycles. It is in the direction opposite that of the deviation generated with the outstretched arm in the Free and Pointing conditions of the current study reported here.

**These results mean that there are two distinct processes evoked by structured visual motion. Arm, head, and torso deviations can be elicited by the onset of visual stimulation prior to experienced self-motion. Compensation for “expected” but absent Coriolis force can be elicited by visual stimulation but only after the perception of self-motion and displacement has occurred. For the same direction of visual stimulation, the two effects are opposite in direction. Although both effects result from active motor control, an important difference is that the pseudo-Coriolis effect requires an active movement and the Deviation Effects do not, making the latter more likely to occur.** One important requirement that is common to both the pseudo-Coriolis effect and the Deviation Effects described here is that the visual environment must include visual cues that are capable of inducing self-rotation. If rotational cues are not present in a visual flow field, our initial test indicates that self-rotation will not normally be perceived and the Deviation Effects described here will not occur. For example, in preliminary tests, we were unable to produce the Deviation Effects with a strictly linear flow field.

Returning to our mention in the Introduction of the first lunar landing, Armstrong was exposed to a strong variegated dust stream moving leftward that affected his ability to control the descent of the lunar module. An excerpt from the 1969 Technical Debrief (Jones, 1995) makes this clear:

Armstrong – “It was a little bit like landing an airplane when there’s a real thin layer of ground fog, and you can see things through the fog. However, all this fog was moving at a great rate which was a little bit confusing.”
Aldrin – “I would think that it would be natural looking out the left window and seeing this moving this way that you would get the impression of moving to the right, and you counteract by going to the left, which is how we touched down.”

From this description, it is clear that the dust stream interfered with their view of the lunar landscape and the visual flow field induced self-motion to the right which likely resulted in control inputs to the left. Taken together these are both disruptions that would favor erroneous control inputs to the left. At the same time within a few seconds of onset, this visual flow field to the left may have induced the arm Deviation Effects described here, which would also produce arm control deviations to the left. Our experimental results predict the error control pattern that Armstrong made. They show that visual motion can elicit unintended deviations of the goal-directed arm both for rotational visual flow alone and for experimental self-rotation and displacement. Appendix 3 below presents evidence that these effects are strong enough to bias joystick control. Our findings from the Pointing conditions with and without the rod indicate that leftward arm deviation excesses would be expected and that their direction corresponds with Armstrong’s actual performance.

Our goal in emphasizing these findings is to call attention to the potential consequences of using HMD displays that contain large visual flow fields with rotation inducing cues and/or involve whole-field visual motion. They can induce unintended arm, head, and torso movements and a sense of both self-motion and displacement, all of which could in turn cause accidents. We will go into detail in a future publication but we have found that the results described above for visually induced illusory self-rotation and displacement also occur for either direction of apparent self-rotation, for one or both arms when standing, for the outstretched arms when seated on a stool with feet on the floor, for the outstretched leg or legs during seated conditions, and for the head and torso as well. The impact of the visual field flow rate, area, and structure on the magnitude of the Deviation Effects described here remains to be fully determined. Our experiments were performed with an HMD virtual environment, but there is no reason to expect that these Deviation Effects would not occur in analogous real-world situations as indicated by the lunar landing example. The conditions that could produce dangerous Deviation Effects include those in which:

1. The operator is freely standing or there is no restraint on torso rotation, or
2. There is no support of the arm or hand to help the operator detect arm/hand drift, and
3. There are no other usable cues to detect arm, torso or head deviations, and
4. The operator is exposed to a large area rotation inducing visual flow field, and
5. There is a temporary or extended degradation or loss of the feedback error signal in the real world or HMD.

## Supporting information

Supplementary - Analysis of Balance

Supplemental Table 2 - Postural Balance SDF Statistics

## Appendix 1 Individual Differences

Subjects showed extremely large azimuthal arm deviations in the CW direction during the Free arm condition as well as head and torso deviations. There was a large variance in the rate of arm deviation during the Free condition, see Figures 2 and 3. Subjects’ awareness of their arm deviation varied considerably. A few were unaware that their arm had deviated, while others realized that their arm was deviating but underestimated the magnitude of its azimuthal angle. Of those subjects who perceived their arm physically deviating, some were surprised and could not believe that their “arm was moving on its own”, and some found the non-volitional arm motion funny and laughed in surprise. As seen in Figure 2, the average maximal arm deviation was between 50 to 60 degrees (though the maximum individual subject spatial deviation was over 150 ° because of the torso rotation), and at an average velocity of approximately 1°/s. The torso and head deviated less, about one-third to two-thirds as much as the arm, and their average angular speeds of deviation, respectively, were almost a third and two-thirds of that of the arm.

A more surprising individual difference was that these Deviation Effects do not occur in the same direction for all subjects even though the induced motion illusion was in the same direction (CCW) for all participants. Of the 12 subjects in this experiment, 11 of them showed arm movement deviations in the direction opposite that of induced self-motion, for both their Free and Pointing conditions. However, one subject showed deviations in the same direction as induced self-motion. All of his deviations were mirror images of the other 11 subjects. In addition to the subjects in this experiment, we tested 12 additional people in preliminary trials or demonstrations of these effects, and one of them also showed mirror image responses. Both of these subjects were righthanded. The one person we tested who was left-handed showed a normal pattern, so this difference does not appear to be related to handedness.

During post-experiment debriefings, two of the subjects reported that holding their eyes fully deviated to the right for 75 seconds in the control condition was uncomfortable, and they may have involuntarily moved their heads slightly to the right to reduce discomfort. If that were the cause of the head deviation, then it would not apply to our experimental conditions, where the subjects never had to uncomfortably voluntarily move their eyes to extremes. In any case, this control condition did show a head deviation with respect to the torso; however, as the re torso section of Table 1 shows, no such effect occurred during the actual experiment. That no head re torso effect was shown in the experiment indicates that it occurred in the eye deviation control condition due to voluntary eye deviation.

## Appendix 2 Relation to Previous Research

Previc and Neel (1995) examined the postural and manual control of subjects viewing scenes consisting of red squares against a dark background, with the scene rotating in roll at 25°/s. Four different size/eccentricity display conditions were carried out for subjects standing on a force plate viewing the different displays while fixating a stationary white target. On a separate day the subjects were seated in a “flight seat” and used left-right movements of a joystick with a light grip of their right hand at the central point of the joystick to maintain the center portion of the display horizontal while being exposed to surround motion. In the posture study, motion in the periphery of the visual field was most effective and caused a .64 cm shift in the center of pressure (COP). The center condition was least effective, biasing the COP by only .15 cm. Our force plate results are consistent with these findings.

The manual task showed substantial biases for full field rotational motion. The visual horizontal setting of the display was 15.2° in the direction of surround motion. This is the orientation of the center display when it appeared to be horizontal. Previc (1992) likened the bias in setting the center display in the manual task to a potential analog of the “Giant Hand Effect” reported by pilots. Some subjects in fact reported that the stick was not responding to their attempts to center the display at horizontal. When subjects used the stick to track an unstable altitude display, they did not sense any disruption of their stick movements, and did not feel a giant hand effect. The displays used in these early studies were state of the art at the time and followed the pioneering studies of Brandt and Dichgans (Dichgans and Brandt, 1978). These arrays readily induce the illusion of “vection”, apparent self-motion. The Previc studies involved vection ratings but they did not mention whether their subjects experienced both self-motion and displacement. With roll stimulation, a peak self-displacement can occur with the subject experiencing roll deviation that peaks with motion still being experienced (Dichgans et al., 1972)

Dvorkin, Kenyon, and Keshner (2009) studied the influence of visual roll motion on the reaching movements of subjects performing double-step reaching tasks. The subjects were exposed to a complex 3D environment that could be rolled at 130°/sec. The scene when rotated began its motion at the onset of a target presentation position that was maintained constant for 2 s in single step trials, in dual step trials the first target appeared for 50, 200, or 500 ms and then a second target appeared for 2 s. Their experiment revealed a disruption of reach kinematics both temporal and spatial. There was a displacement of the final position of the subject’s hand in the direction of the visual roll of the visual scene that was consistent with “---interference with the ability to simultaneously process two consecutive stimuli” (Dvorkin et al, 2009, pg. 95).

The exposure times to the visual roll of the display were too short to induce apparent self-motion. Subjects received 20 practice trials before the start of data collection. In the Coriolis paradigms for studying reaching movements to targets (Cohn et al., 2000, Lackner and DiZio, 2000) practice trials were not given. After 20 perturbed reaches adaptation was approximately 80% complete. It is possible that the effects observed by Dvorkin, et al, 2009 would be even larger if practice trials were not given.

Bringoux and colleagues explored whether an object held in a pincer grip at a seated subject’s lap would show systematic anticipatory grip compensations for visually induced oscillatory vection in the absence of real motion (Bringoux et al., 2012, White et al., 2020). Such anticipatory adjustments of grip force rapidly develop during actual exposure to up-down oscillatory motion. A visually induced effect on precision grip was not elicited despite their ability both to record precision grip with great accuracy and to induce compelling apparent up-down oscillation in their subjects. Our finding that subjects during experienced self-rotation and displacement did not sense the rod as exerting a centrifugal force on their hand is in accord with that of Bringoux and colleagues (Bringoux et al., 2012).

## Appendix 3 Effects on Control of Joystick

To determine whether the arm deviations that we evoked in our Free and Pointing conditions could influence the control of a joystick, we conducted additional conditions. Twelve subjects participated in two joystick conditions. One was with a joystick that, with an outstretched arm, they attempted to set to the midpoint of its ± 20° range of roll motion between mechanical stops. Subjects held it at its top with their fingers attempting to center it while they were exposed to the rotating visual scene. The joystick was centered before the start of a trial after subjects had moved it through its range of motion. All subjects experienced illusory CCW self-motion within 10 s of exposure to the CW rotating visual scene. Within 5 s of self-motion subjects began to move the joystick rightward, with it typically “pegging” at its 20° rightward limit within 20 s of trial onset and remaining pegged until the end of the 75 s duration trial. Most subjects expressed surprise when they felt the joystick hit the right stop. In the other condition, subjects used a heavily damped joystick, where less deviation occurred as subjects detected its resistance to their involuntary arm deviation. In preliminary tests, these joystick deviations did not occur when a linear visual flow field was used; the visual scene had to include self-rotation inducing cues. These and related observations will be described in a paper in preparation that includes extended observations on the rod as a tool that can capture and extend the base of experienced self-rotation.

One way to isolate or at least minimize Deviation Effects in the joystick situation would be to provide a hand or wrist support close to the joystick. The friction between the support and the hand would tend to prevent the arm from drifting, and help isolate the joystick control from the arm Deviation Effect. This would be true unless Deviation Effects operate all away down to the level of the hand and fingers. Initial tests indicate that a wrist support does help to isolate joystick operation from arm deviation-like effects.

## Appendix 4 Why Do Pointing Errors re Space Occur?

Exposure to a continuously rotating textured environment would have been highly improbable during human evolution and humans possibly never evolved to correct for passive body deviations resulting from illusions of self-rotation. However, it is more likely that the motor control system does have access to arm re space information, but cannot or does not use it because of the self-rotation illusion. The subject’s feet were stationary on the force plate and the force plate was fixed in relation to the briefly presented tactile target in the Pointing conditions. Therefore, the sensors in the subject’s muscles and joints from their feet through the legs, torso, and arm should enable the subject to use that information to point accurately in space as well as with respect to their torso. However, because the subject has the illusion that they are rotating with respect to the fixed tactile target, the motor control system may ignore that information.

As a preliminary test of this hypothesis we tested four subjects informally in a pointing task in which they again had to point to a tactually defined location in space while viewing either a rotating or stationary display in the HMD. At the same time their upper torsos were physically rotated CW below the velocity detection threshold (less than 1° per second). When viewing the rotating display in the HMD, their pointing remained relatively fixed with respect to their torso and deviated from the target location in space as the torso turned. This is just what we saw in the formal experiment. However, when viewing the stationary display, the subjects pointed much more accurately to the fixed location in space. This indicates that the information to point accurately in space is available to the motor control system, but the illusion of self-rotation results in the motor control system’s failure to use that information. However, the illusion of self-rotation does not interfere with the ability of the motor control system to accurately locate the arm with respect to the torso. The important difference here is probably that the torso and arm are being perceived as rotating together, but because of the self-rotation illusion, the feet are perceived as rotating with respect to the fixed target location in space. This is an unusual example of a conscious illusion interfering with a subconscious motor control process.

## Author Contributions

JRL first noticed the arm deviation effect, was involved in planning the experiment, and took part in the experiment’s execution. AM was the primary experimenter, recruiting subjects and jointly running the experiment. JV contributed to the experimental design and conduct of the experiment, and was the primary statistical consultant. SP developed the virtual reality environment. AB participated in conducting the experiment, coded the data analysis scripts in MATLAB, prepared the figures, and did the statistical analysis in SPSS. All authors contributed to preparing the manuscript.

## Dedication

This paper is dedicated to Dr. Angus Rupert. He told JRL about the control problem experienced by Neil Armstrong during the moon landing and called JRL’s attention to the remarks by Armstrong and Aldrin that are cited in the text. During pilot tests before the execution of the experiments described in the current article, JRL recognized their significance for the situation Armstrong faced.

## Acknowledgments

We thank Alberto Pierobon for technical hardware support and Lee Picard for editorial support.

