## Supplementary - Analysis of Balance for "Visually Induced Involuntary Movements"

A Stabilogram Diffusion Function (SDF) analysis of the force plate results and their relevance for the present findings are presented in this online supplementary material.

***Methods***

During experimental trials, subjects stood on a dual force plate (AMTI Accusway$^{TM}$) that allowed separate monitoring of the forces and torques exerted by each foot. Sampling rate was 200 Hz. Fifteen dependent variables were derived from the force plate data.

The force plate variables were divided into three groups:

1. Anterior-Posterior Center of Pressure ($COP_{AP}$), which characterized how the subject's COP moved in the fore-aft plane,
2. Medial Lateral Center of Pressure ($COP_{ML}$), which characterized the subject's COP motion in the right-left plane, and
3. Weight Fraction (WF), which characterized how the subject distributed their weight between their two feet over time.

Center of Pressure (COP) trajectories in quiet passive stance are dominated by stochasticity and metrics such as mean sway amplitude and velocity, are inadequate to describe fully postural sway (Bakshi A et al., 2014;Collins JJ and De Luca CJ, 1993). Consequently, we used the Stabilogram Diffusion Function (SDF), which is sensitive to the stochastic nature of quiet stance. From the force plate measures above, we calculated SDFs, which measure the mean square displacement (MSD = C) of a time-series as a function of temporal window width $\Delta\tau = t_2 - t_1$. Thus, $C(t_2 - t_1) = \langle[\theta(t_2) - \theta(t_1)]^2\rangle$, where $\theta$ represents time-series variables (for example,

$COP_{AP}$, $COP_{ML}$, and WF as a function of time t). The SDF characterizes both the stochastic and deterministic features of postural sway (De Luca C and Collins J, 1995). Each SDF trace is further characterized by the following metrics, which generate five additional dependent variables used in the statistics:

1. SDF Area—The total Area under each of the SDF traces.
2. Hurst Exponent (H) – Exponent 2H determined by the slope of log(C) plotted against log($\Delta\tau$) at the shortest time scale ($\Delta\tau \rightarrow 0$), where $C=\langle\Delta\theta^2\rangle = D_{eff}\,\Delta\tau^{2H}$.
3. Critical Point (t*) – the time at which the exponent H =0.5.
4. Effective Diffusion Coefficient ($D_{eff}$) – The upfront constant that measures the stochastic activity on average.
5. SDF Halfway Magnitude ($t_{1/2}$)– The mean square displacement at the halfway value of the largest temporal bin analyzed, at time bin width $\Delta\tau$=1 s.

We performed three 3-way MANOVAs, one for each group of variables in the three areas of interest. The MANOVAs tested for main effects of the independent variables - 1. Epoch with three levels (Pre, Initial, Remainder), 2. Task with two levels (Free, Pointing), and 3. Tool with two levels (rod, no-rod). We also tested for significance of all possible interactions among the three independent variables. Table 2 shows the full testing structure in detail.

### Force Plate Results

The statistical results for the Force Plate Data are shown in Table 2. The degrees of freedom for the multivariate effects tests involving Epoch are $F(5,19)$, and for those just involving Task and Tool are $F(5,7)$. The degrees of freedom for the univariate ANOVAs are all $F(2,22)$ and for the follow up post hoc contrasts involving Epoch are F(1,11). The significance levels and Partial Eta Squared (□$^2$) effect sizes are shown in Table 2. Figures 5 shows the Stabilogram Diffusion Functions (SDF) for the AP components of the COP fluctuations. The statistical details of the five parameters characterizing the SDF are shown in the AP COP MANOVA section of Table 2. The only significant multivariate effect in the AP COP MANOVA is for the Epoch independent variable ($\Theta$=2.9, F(5,19)=10.9, p <.001 , $\eta^2$=.74), there were no Task or Tool effects. Visual inspection of Figure 5 suggests that this was at least in part due to the higher mean square displacement (MSD) in the Remainder Epoch versus Initial and/or Pre. This conclusion is supported by the significant SDF Area univariate ANOVA. In addition to the SDF Area (p<.03, $\eta^2$=.28), the Hurst Exponent (p<001, $\eta^2$=.69) and halfway magnitude (p<.03, $\eta^2$=.28) were significantly affected by Epoch. The post hocs show that for the Hurst Exponent both the Pre vs Initial and Initial vs Remainder differences were statistically significant. For the SDF Halfway Magnitude values only a linear combination of the effects in the three levels of Epoch can account for the significance.

Figure 6 shows the SDF for the ML component of the COP fluctuations. Statistical details for the five dependent variables derived from the SDF are shown in the ML COP MANOVA section of Table 2. The only significant multivariate effect is for Epoch ($\Theta$=1.3, F(5,19)=4.9, p<.01, $\eta^2$=.56), no Free versus Pointing differences were found, nor were any Tool effects found. The pattern of significant univariate ANOVAs for these dependent variables is similar to that for the AP COP but for the ML COP, the Diffusion Coefficient is also significantly affected by Epoch (Area p<.04, $\eta^2$=.27; Hurst p<.01, $\eta^2$=.39; Diffusion Coefficient p<.05, $\eta^2$=.28; and Halfway Magnitude p<.02, $\eta^2$=.31). Unlike for AP COP, where the post hoc tests showed there were a mixture of effects from the different levels of Epoch causing the significant results, in the ML case the post hoc tests show that, without exception, it is the difference between the Initial and Remainder levels of Epoch that is responsible for the statistically significant results.

Figure 7 shows the SDF for Weight Fraction (WF), and the WF MANOVA section of Table 2 has the statistical details. For the WF SDF, it is difficult by visual inspection to discern any consistent pattern of differences across the conditions. Note that the left and right feet have the same SDF traces since $WF_{RT} + WF_{LT} = 1$, so $\Delta WF_{RT} = \Delta WF_{LT}$, and the MSD of fluctuations will be identical for both feet. As was the case for the two COP MANOVAs, the only significant multivariate main effect is for Epoch ($\Theta$=1.2, $F(5,19)$=4.6, $p$<.01, $\eta^2$=.55). There are no significant Task or Tool effects. But unlike in the COP MANOVAs, the significance only results from the Hurst Exponent ANOVA, which is highly significant at $p$=.004, $\eta^2$=.40. Since this exponent is computed at a very short-time-scale, it implies the WF SDFs show differences across Epoch only at very short time-scales. The post hoc contrasts show that this effect results from differences between the Initial and Remainder levels of Epoch. There is also a significant multivariate Epoch x Tool interaction ($\Theta$=.9, $F(5,19)$=3.5, $p$<.03, $\eta^2$=.48) that results from significant Hurst Exponent and Critical Point dependent variable ANOVAs. The Hurst Exponent post hocs show that the significant difference is due primarily to the difference between the Pre and Initial levels of Epoch. This is the only case where a significant difference between these two Epoch levels was observed. For the Critical Point dependent variable, the post hocs show that the differences between the Pre and Initial levels of Epoch, as well as those between Initial and Remainder are significant. Unlike for the AHT data, for the force plate data the only significant independent variable is Epoch. There is no effect of Task or Tool; in fact, none of the balance metrics show

Task or Tool dependency as the AHT measures did. Unlike for the AHT data, the only significant interaction is between Epoch and Tool. Since the rod added additional weight on the right hand, the effect of its presence affected the WF fluctuations more when the arm and rod became laterally extended than when extended forward.

Overall, among the balance metrics from the force plate, the dependent measures derived from the SDFs of $COP_{AP}$, $COP_{ML}$ and WF showed significant effects for Epoch. Are these effects arising due to the visual flow effects or illusory self-motion and displacement, or is it the physical arm deviation influencing posture? The arm deviation variables (angular position, and average velocity during the arm deviation) showed significant effects both due to Epoch (Pre, Initial, Remainder) and the Task (Free vs. Pointing). By contrast, among the postural balance metrics from the force plate, the dependent variables derived from the SDFs of COP and WF showed significant effects due to Epoch but not Task. **This supports that effects on postural balance originate from visual motion and are not confounded by the arm being laterally deviated more in the free conditions versus less so in the pointing conditions.**

**Discussion**

Subjects in any small interval of time were mostly standing in a quiet stance. Quiet stance COP traces are dominated by stochasticity, which is why we used the SDF analysis to characterize postural changes. The COP component in the forward-backward direction showed significant changes after the onset of self-motion perception. Specifically, the short-time-scale power law exponent and the critical point (location in the temporal window axis where the SDF traces inflect from concave to convex, or Brownian to corrective motion) showed differences between Pre and during self-motion intervals.

Previc and colleagues were the first to raise the important question whether moving visual arrays could influence postural and joystick control. They wondered whether a "giant hand" type illusion could be induced visually that would be analogous to that experienced in flight when a pilot feels that his or her commands to the aircraft are not being followed because it feels as if a giant hand were tilting the aircraft in roll. In a series of early experiments, they measured postural sway and stick control under conditions of various patterns of moving visual stimulation (Previc FH and Donnelly M, 1993;Previc FH and Ercoline WR, 2004;Previc FH et al., 1993;Previc FH and Mullen TJ, 1991;Previc FH and Neel RL, 1995). They also looked at the relationship between the onset of visual motion and that of apparent self-motion, "vection", and postural change. In one study, subjects standing on a force plate viewed a visual scene projected on a screen subtending 93° horizontally and 81° vertically (Previc FH and Mullen TJ, 1991). The visual scene consisting of 100 small white squares against a dark background, could be rotated at 25°/sec in clockwise or counterclockwise roll. After 10 sec of viewing the scene static, it was then set in motion for 50 sec. Postural changes to stimulus motion occurred in less than 5 sec, and vection occurred at just over 7 sec. Most subjects first postural change was a lateral shift of COP about 1.5 cm leftward for CCW visual motion and rightward for CW roll motion stimulation. Despite the slight lag in vection onset to force plate changes, ratings of vection magnitude and onset showed a strong correlation with force plate changes reflecting postural shifts. Our SDF analysis shows postural changes that are in accord with Previc and Mullen's observations.

**Figure Captions**

**Figure 5**. The Stabilogram Diffusion Functions (SDFs, see method section for details) for the fluctuations in the anterior-posterior (AP) component of the Center of Pressure (CP), "$CP_{AP}$ fluc". The SDF is computed for temporal windows of variable sizes, up to a maximum width of 2 s. The top panels represent the free conditions, bottom panels the pointing conditions. The Left and right panels are for no-rod and rod conditions, respectively.

**Figure 6**. SDF for fluctuation in the CP's mediolateral (ML) component, i.e., $CP_{ML}$ fluctuations.

**Figure 7**. SDF of the weight fraction (WF) fluctuations under either foot (as noted in text, WFs under both feet will give precisely the same traces).


Bakshi A, DiZio P, Lackner JR (2014), Statistical analysis of quiet stance sway in 2-D. Experimental brain research 232:1095-1108.

Collins JJ, De Luca CJ (1993), Open-loop and closed-loop control of posture: a random-walk analysis of center-of-pressure trajectories. Experimental brain research 95:308-318.

De Luca C, Collins J (1995), The effects of visual input on open-loop and closed-loop postural control mechanisms. Experimental brain research 103:151-163.

Previc FH, Donnelly M (1993), The effects of visual depth and eccentricity on manual bias, induced motion, and vection. Perception 22:929-945.

Previc FH, Ercoline WR (2004) Spatial disorientation in aviation. Aiaa.

Previc FH, Kenyon RV, Boer ER, Johnson BH (1993), The effects of background visual roll stimulation on postural and manual control and self-motion perception. Perception & psychophysics 54:93-107.

Previc FH, Mullen TJ (1991), A comparison of the latencies of visually induced postural change and self-motion perception. Journal of Vestibular Research 1:317-323.

Previc FH, Neel RL (1995), The effects of visual surround eccentricity and size on manual and postural control. Journal of Vestibular Research: Equilibrium & Orientation.
