## Supplemental Table 2 - Postural Balance SDF Statistics for "Visually Induced Involuntary Movements"

| | | Force Plate Data MANOVAs | | | | | | |
|---|---|---|---|---|---|---|---|---|
| | | INDEPENDENT VARIABLES | | | | | | |
| | | Epoch | Task | Tool | Epoch x Task | Epoch x Tool | Task x Tool | Epoch x Task x Tool |
| AP COP MANOVA | p-value | **<.001** | .401 | .120 | .249 | .340 | .159 | .589 |
| | effect size | **(.74)** | (.46) | (.65) | (.28) | (.24) | (.62) | (.17) |
| DEPENDENT VARIABLES | | | | | | | | |
| SDF Area | p-value | **.026** | | | | | | |
| | effect size | **(.28)** | | | | | | |
| Hurst Exponent | p-value | **<.001** | | | | | | |
| | effect size | **(.69)** | | | | | | |
| Critical Point | p-value | .055 | | | | | | |
| | effect size | (.26) | | | | | | |
| Diffusion Coef. | p-value | .096 | | | | | | |
| | effect size | (.19) | | | | | | |
| SDF Halfway Mag. | p-value | **.026** | | | | | | |
| | effect size | **(.28)** | | | | | | |
| ML COP MANOVA | p-value | **.005** | .132 | .632 | .189 | .071 | .347 | .224 |
| | effect size | **(.56)** | (.64) | (.34) | (.31) | (.39) | (.49) | (.29) |
| DEPENDENT VARIABLES | | | | | | | | |
| SDF Area | p-value | **.032** | | | .049[2] | | | |
| | effect size | **(.27)** | | | (.24) | | | |
| Hurst Exponent | p-value | **.004** | | | | | | |
| | effect size | **(.39)** | | | | | | |
| Critical Point | p-value | .465 | | | | | | |
| | effect size | (.06) | | | | | | |
| Diffusion Coef. | p-value | **.048** | | | .031[2] | | | |
| | effect size | **(.28)** | | | (.27) | | | |
| SDF Halfway Mag. | p-value | **.017** | | | | | | |
| | effect size | **(.31)** | | | | | | |
| WF MANOVA | p-value | **.006** | .115 | .586 | .230 | **.020** | .170 | .117 |
| | effect size | **(.55)** | (.66) | (.36) | (.29) | **(.48)** | (.61) | (.35) |
| DEPENDENT VARIABLES | | | | | | | | |
| SDF Area | p-value | .145 | | | .046[2] | .69 | | |
| | effect size | (.16) | | | (.244) | (.02) | | |
| Hurst Exponent | p-value | **.004** | | | | **.009** | | |
| | effect size | **(.40)** | | | | **(.35)** | | |
| Critical Point | p-value | .41 | | | | **.043** | | |
| | effect size | (.07) | | | | **(.25)** | | |
| Diffusion Coef. | p-value | .113 | | | .028[2] | .87 | | |
| | effect size | (.18) | | | (.28) | (.01) | | |
| SDF Halfway Mag. | p-value | .070 | | | | .90 | | |
| | effect size | (.22) | | | | (.001) | | |
| | | Epoch | Task | Tool | Epoch x Task | Epoch x Tool | Task x Tool | Epoch x Task x Tool |

| | AP COP Epoch Post Hocs | | ML COP Epoch Post Hocs | | | POST HOC CONTRASTS FOR WF MANOVA: Epoch | | Epoch X Tool | |
|---|---|---|---|---|---|---|---|---|---|
| | pre vs initial | initial vs remainder | pre vs initial | initial vs remainder | | pre vs initial | initial vs remainder | pre vs initial | initial vs remainder |
| SDF Area | .342 | .091 | .880 | **.027** | SDF Area | | | | |
| | (.08) | (.24) | (.00) | **(.37)** | | | | | |
| Hurst Exp | **.040** | **<.001** | .965 | **.007** | Hurst Exp | .480 | **.003** | **.017** | .279 |
| | **(.33)** | **(.73)** | (.00) | **(.50)** | | (.05) | **(.58)** | **(.42)** | (.11) |
| Crit Point | | | | | Crit Point | | | **.038** | **.008** |
| | | | | | | | | **(.34)** | **(.49)** |
| Diff Coef | | | .976 | **.007** | Diff Coef | | | | |
| | | | (.00) | **(.50)** | | | | | |
| SDF Half | .550 | .067 | .918 | **.018** | SDF Half | | | | |
| | (.03) | (.27) | (.00) | **(.41)** | | | | | |

Note 1: [1] Indicates that the p-value for this dependent variable is included, whether significant or not, even though the MANOVA for this factor was not significant, because it was an a priori prediction.

Note 2: [2] Indicates the univariate p-value for this variable was significant, but this p-value will not be interpreted by us because the MANOVA for this factor was not significant and there was no a priori prediction. These values are only included for future speculation.

**General Rules followed in constructing this table.**

1. The statistical significance level for all entries in this table was set to $p < .05$.
2. For each effect, the p-value is in the first line, and the effect size (eta squared) is within parenthesis in the second line.
3. Results from a priori predictions are included and marked with superscript 1 and printed in **black-bold** if statistically significant or in gray if not.
4. All MANOVA results are included and printed in **black-bold** if significant or in gray if not.
5. All univariate results are included if the corresponding multivariate factor was significant. These univariate results are printed in **black-bold** if significant or in gray if not.
6. All other univariate results are omitted unless significant, in which case they are superscripted 2 and printed in gray .
7. The inclusion of post hoc results for Epoch and its interactions followed analogous rules as in 5 and 6 above.
8. Only those results printed in **black-bold** are interpreted by us in this paper.

**Table 2. Summary of Force Plate SDF Statistical Results**
